## Supplement for "Biomass, abundances, and abundance and geographical range size relationship of birds along a rainforest elevational gradient in Papua New Guinea"

Figure S1. Spearman correlation between *mean abundances* of all bird species recorded during point-counts (PC) and during mist-netting (MN, data from (Sam et al. 2019). The correlation between the data was rather close, with some birds being recorded only during point-counts but not during mist-netting. Typically, these were canopy species like pigeons and doves. A species which was often recorded during point-counts but only rarely to nets was a canopy occupying honeyeater *Melidectes belfordi* (abundances 19.8 in PC vs. 2 in MN).

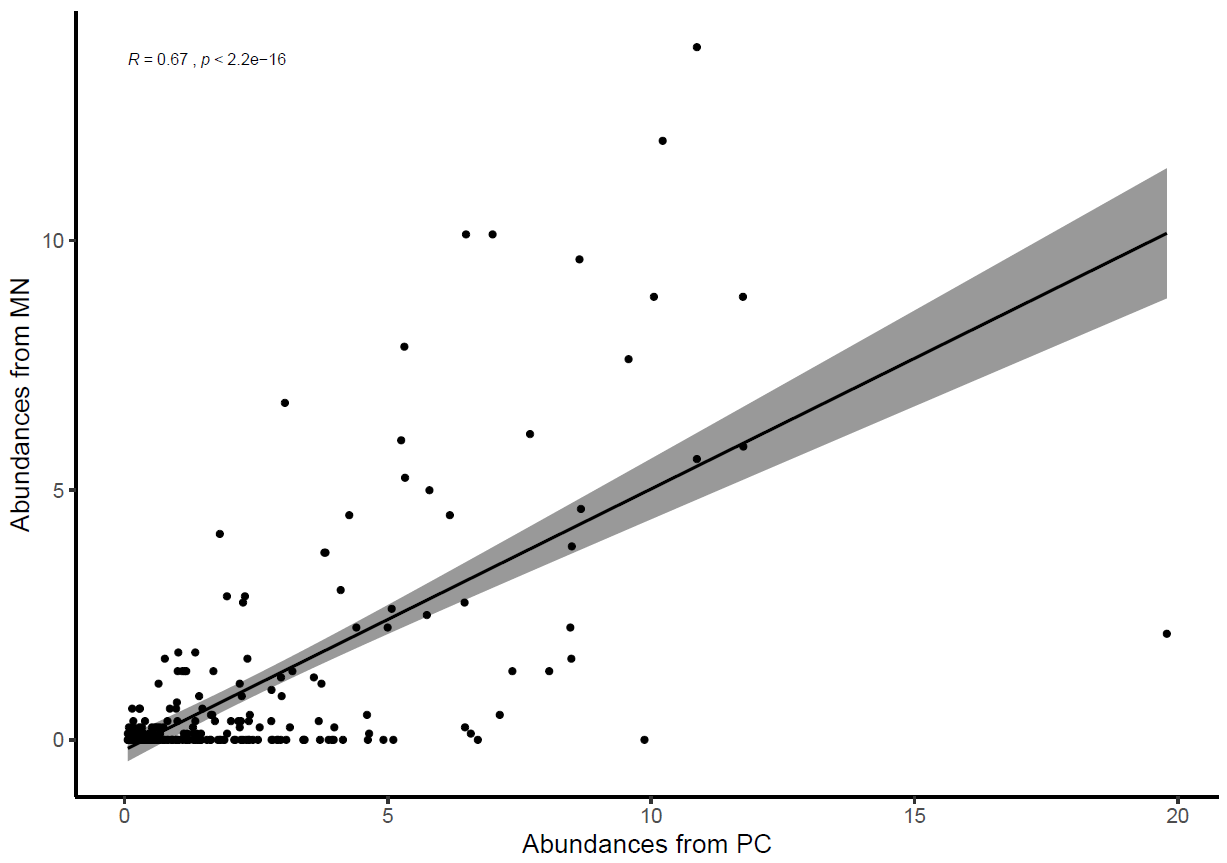

Figure S2. Non-passerine and passerine birds divided into three groups based on the position of their weighted mean point of elevational distribution on Mt. Wilhelm and their *mean abundance* obtained from mist-netting data (data from (Sam et al. 2019) of individual species across elevations (a) and their range sizes in km^2^  (b). Significant differences between the groups of birds are denoted by different letters above the box-plots. Note log scale used on y-axis and different scale of y-axes in part a and b. Lowland group = elevational weighted mean point up to 800m a.s.l., mid group = elevational weighted mean point between 801 and 1600m a.s.l., and montane group = elevational weighted mean point above 1600 m a.s.l. : Kruskal-Wallis test for Passerines (N = 161) (a) χ^2^ = 22.4 , df = 2, N = 161, P < 0.001, (b) χ^2^ = 67.3 , df = 2, N = 161, P < 0.001. Non-passerines (N = 88) (c) χ^2^ = 1.89, df = 2, N = 88, P =0.388 (d) χ^2^ = 19.546, df = 2, N = 88, P < 0.001. For this analysis, weighted mean point and mean abundance was calculated from mist-netting data the same way as from point-count data.

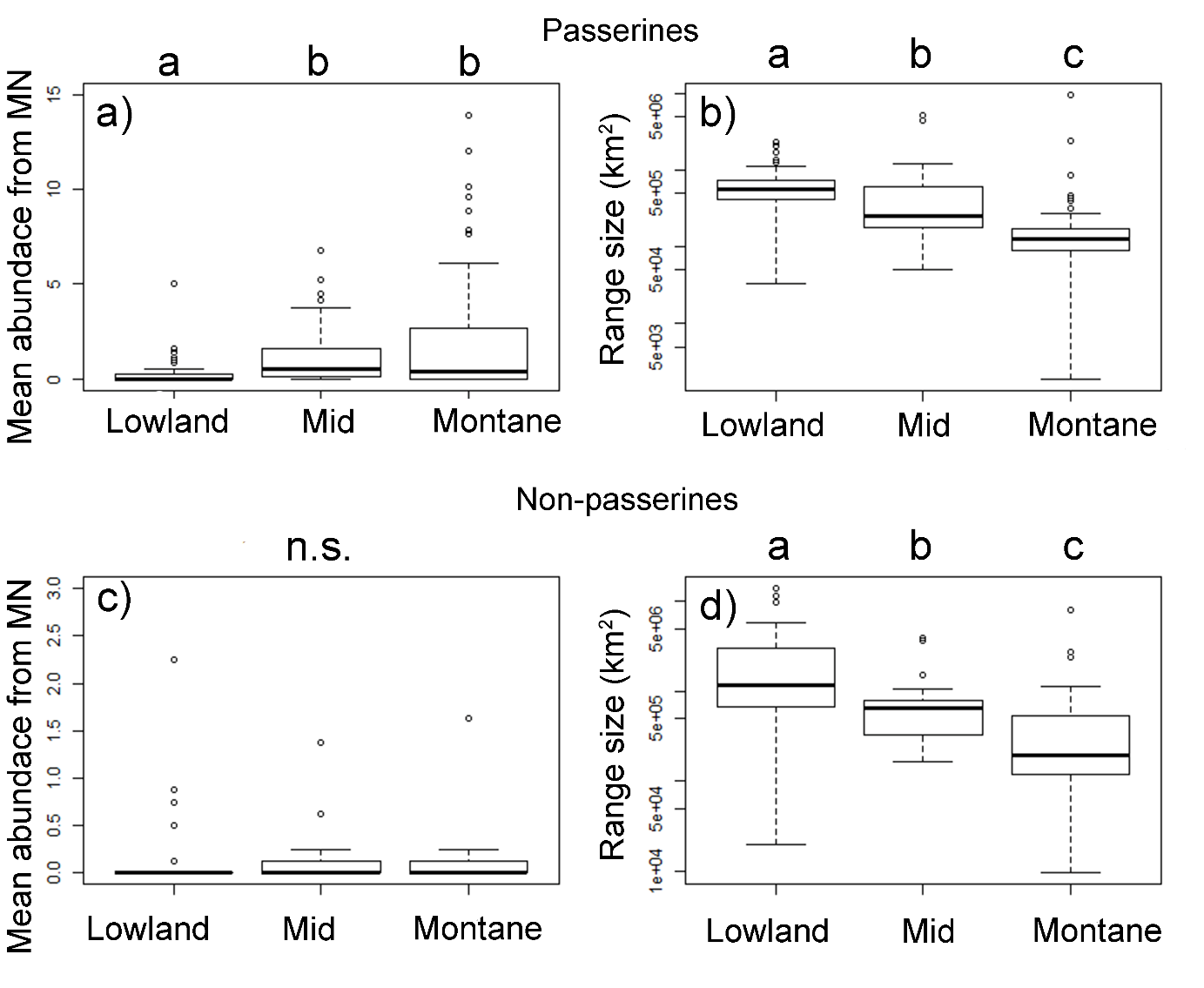

Figure S3. Correlation between *mean elevational abundances* of all bird species recorded during point-counts in wet and dry season (249 species * 8 elevations = > N = 1992). Intercept shows data for passerines only (N = 1288).

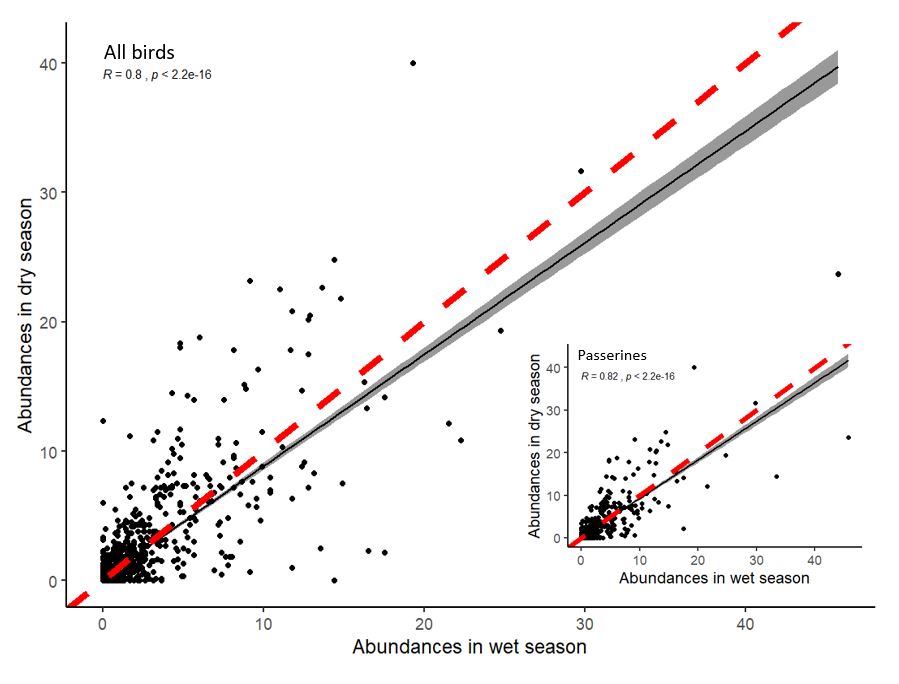

Figure S4. Differences between various measures used in the analyses**.** *Weighted mean point* - the weighted mean is similar to an ordinary arithmetic mean, except that instead of each of the elevational site within the distributional range of a bird species contributing equally to the final average, the elevational points with higher abundance contribute more than others. Note that the point might fall outside of the surveyed sites. *Maximum point* – one out of the eight studied elevational sites where the highest abundace of a bird species was recorded. *Mean elevational abundance* – mean (across 16 points and replicates in time) number of birds recorded per 12.56 ha in 15-minute-long census. For some analyses, the elevational abundance was split into mean elevational abundance in wet season (16 points * 6 replicates in time) and dry season (16 points * 8 replicates in time). *Maximal mean elevational abundance* – the highest mean elevational abundance recorded for the given species along the gradient. *Mean abundace* - mean of mean elevational abudnaces from across the sites where the birds species ocured.

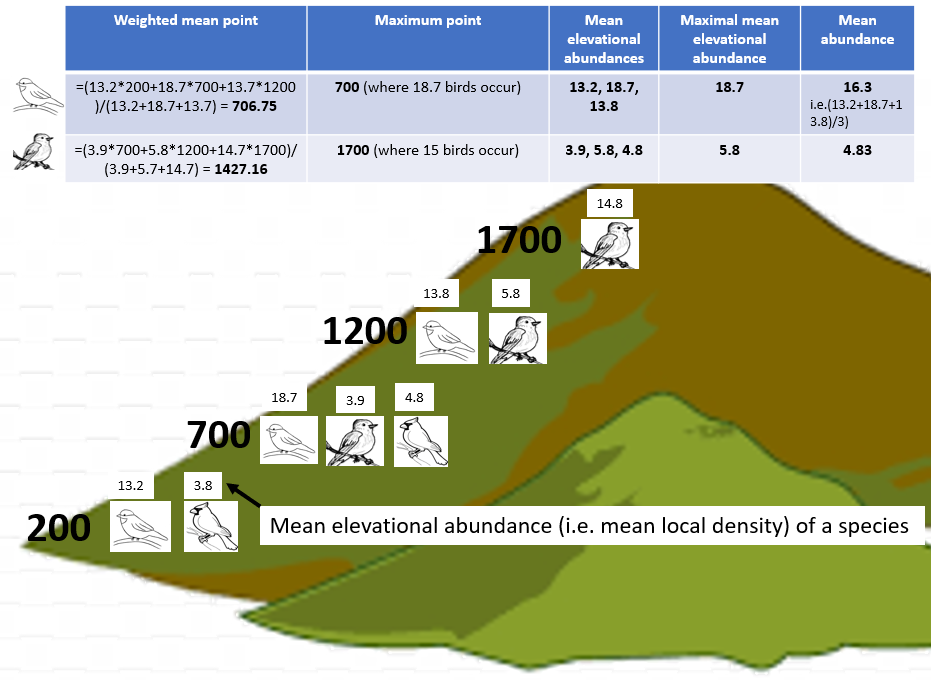

Figure S5. Mean (±SE) number of individuals per passerine and non-passerine bird species occurring in each assemblage at each elevational increment along Mt Wilhelm.

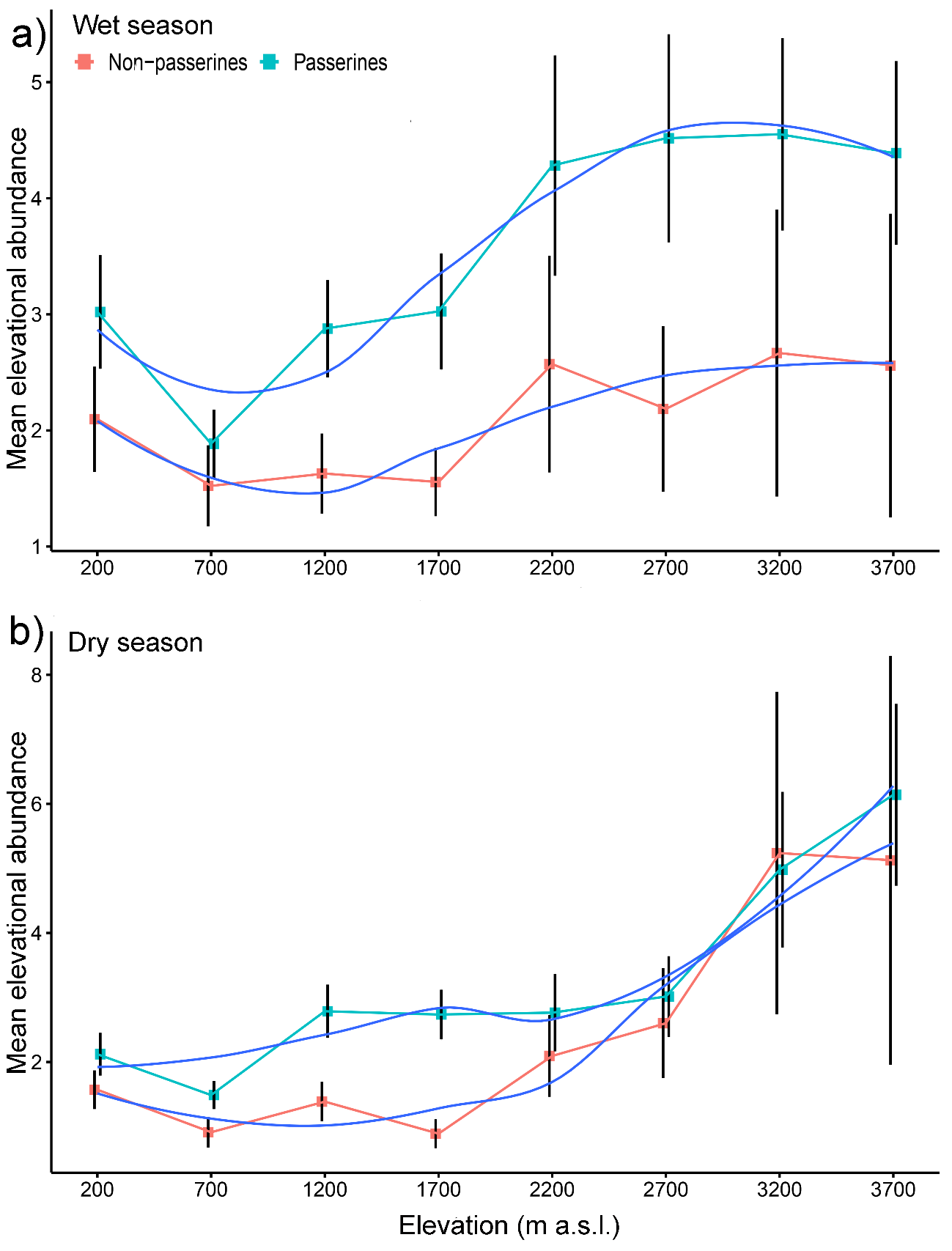

Figure S6. Maximal elevation abundances of passerine and non-passerine bird species with maximal abundances at certain elevation. Bird species with maximal abundances above 1700 m typically had higher maximal abundances than bird species with maxima at lower elevations.

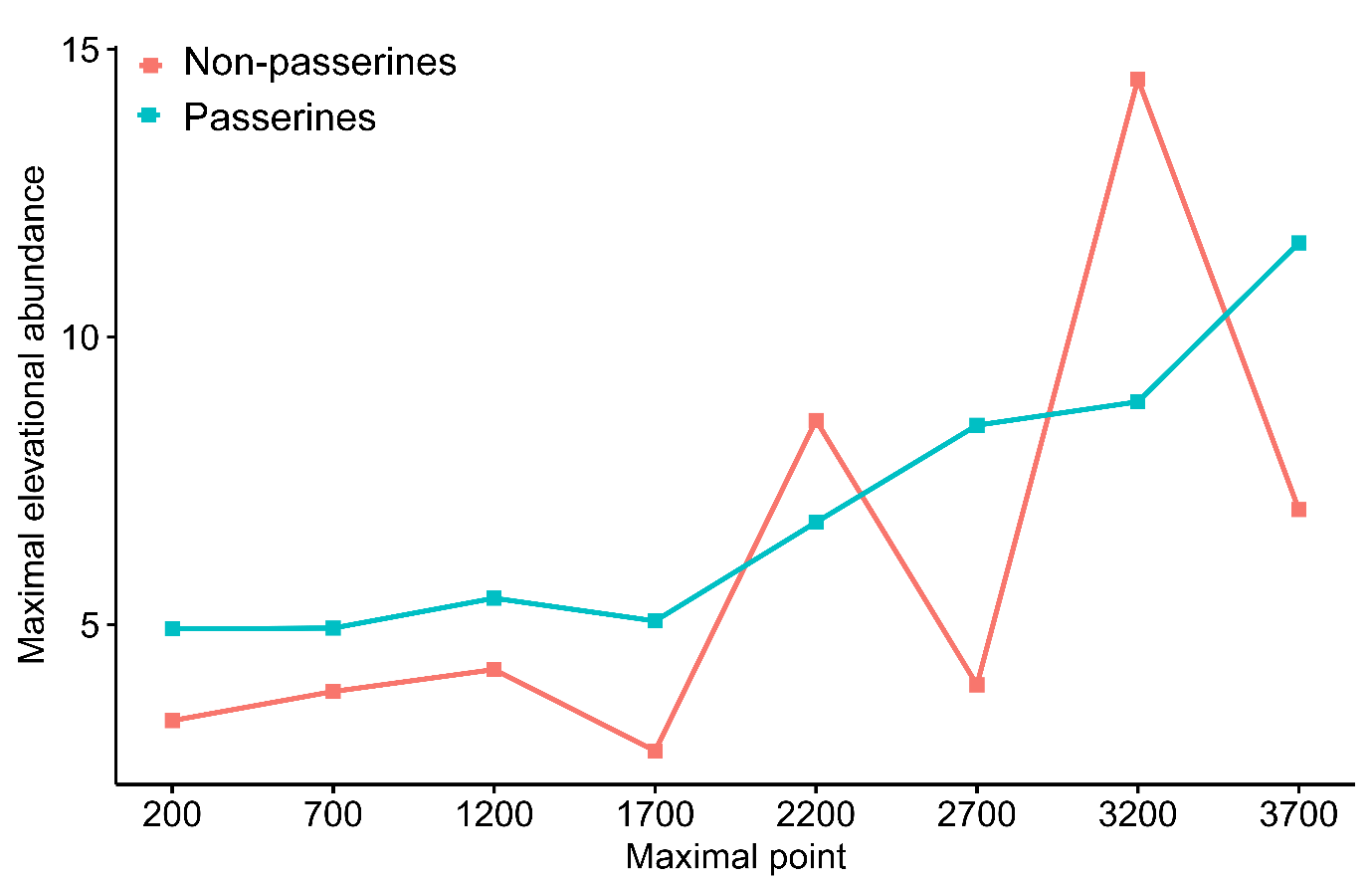

Figure S7. The relationship between the mean abundance of geographical ranges (log transformed) of individual bird species. Only the relationship between mean abundances of all bird species and their ranges was significant (black line, F_1,248_ = 8.22, P = 0.004). After subsampling into passerine and non-passerine birds, the trends remained negative, albeit non-significant, for passerines (F_1,159_ = 1.17, P = 0.28) and non-passerines (F_1,86_ = 2.6, P = 0.10) separately.

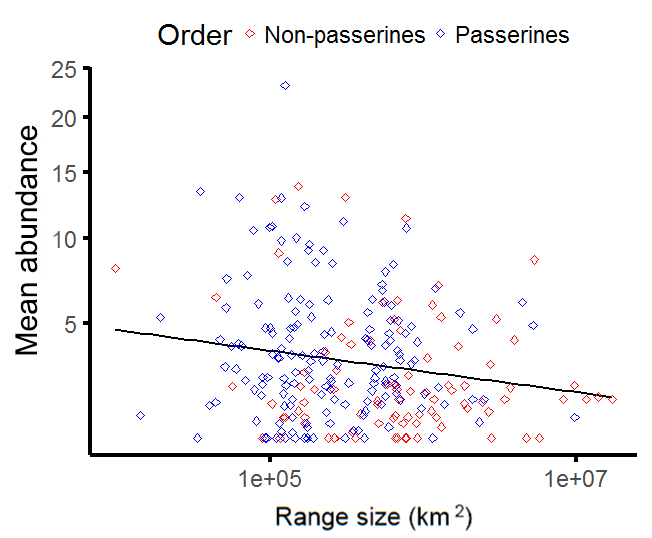

Figure S8. Abundance-range size relationship of three groups of passerine (black dashed lines) and non-passerine (red lines) bird species. (a) species with weighted mean point below 800 m a.s.l. (b) species with weighted mean point between 800 and 1600 m a.s.l. (c) species with weighted mean point above 1600 m a.s.l. Trends are depicted by regression lines fitted by the ordinary least squares method. Note log scale used on x-axes and square root transformation on y-axes. The insets depict the patterns we expected for particular species groups based on range size limitations and increasing abundance towards higher elevations

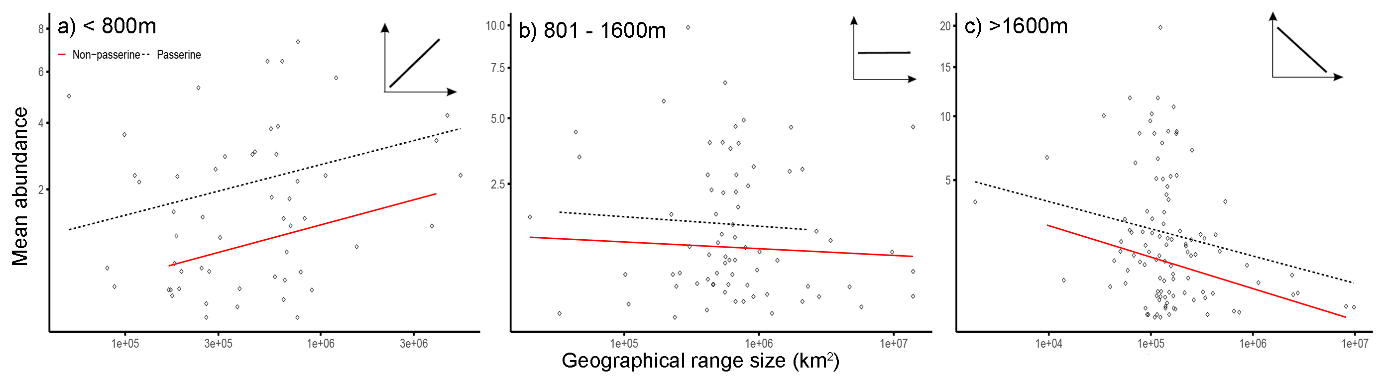

Figure S9. Passerine (a ,b) and non-passerine (c, d) birds divided into three groups based on the position of their weighted mean point of elevational distribution on Mt. Wilhelm, and their *mean abundances* in wet (a, c) and dry season (b, d). Kruskal-Wallis - passerines in dry season (a) χ^2^ = 5.5, df = 2, N = 161, P < 0.05; in wet season (b) χ^2^ = 17.3, df = 2, N = 161, P < 0.001; non-passerines in dry season (c) χ^2^ = 1.9, df = 2, N = 88, P = 0.377; in wet season (d) χ^2^ = 0.5, df = 2, N = 88, P =0.773. Significant differences between the groups of birds are denoted by different letters above the box-plots. Lowland group = elevational weighted mean point up to 800m a.s.l., mid group = elevational weighted mean point between 801 and 1600m a.s.l., and montane group = elevational weighted mean point above 1600 m a.s.l.

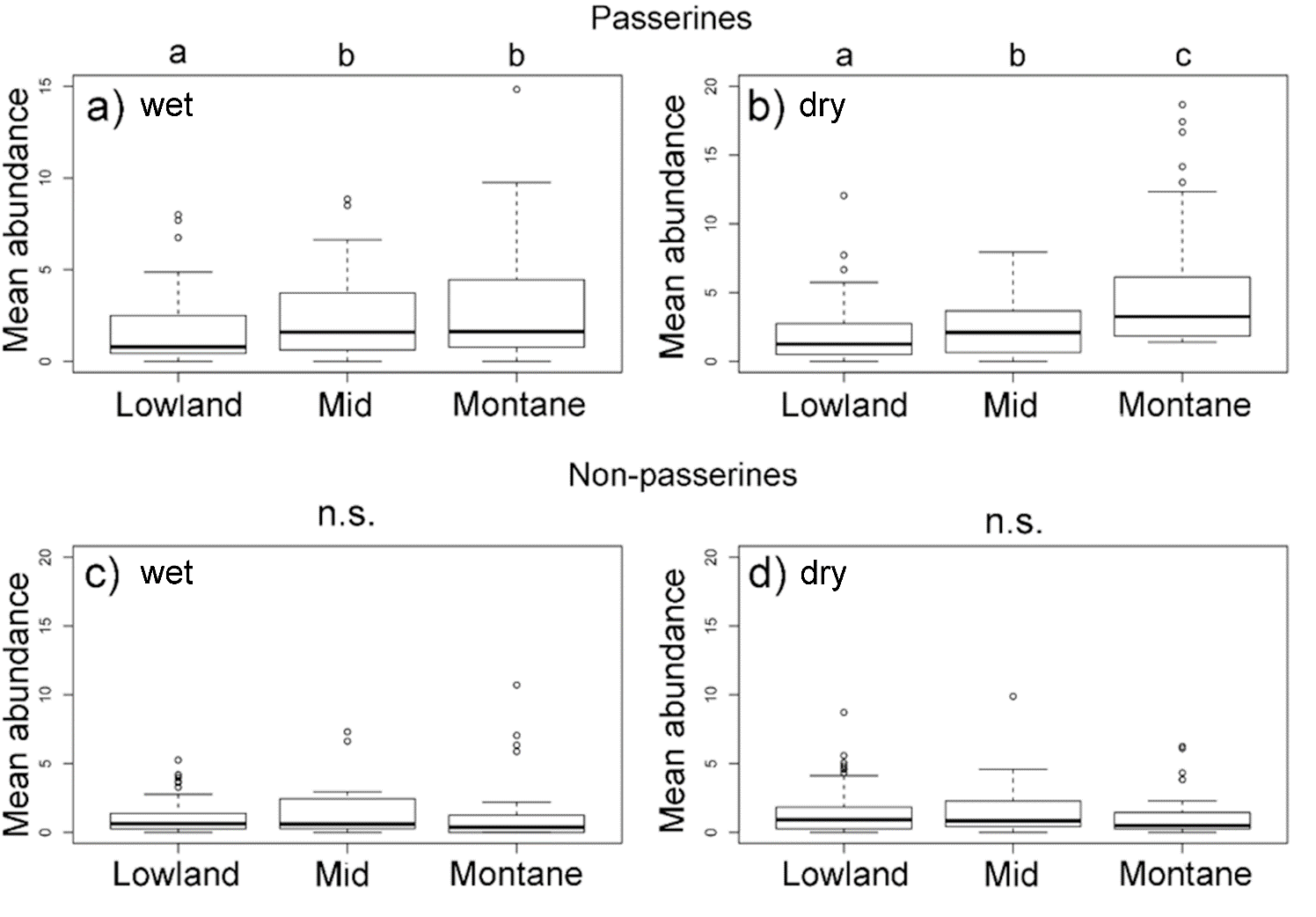

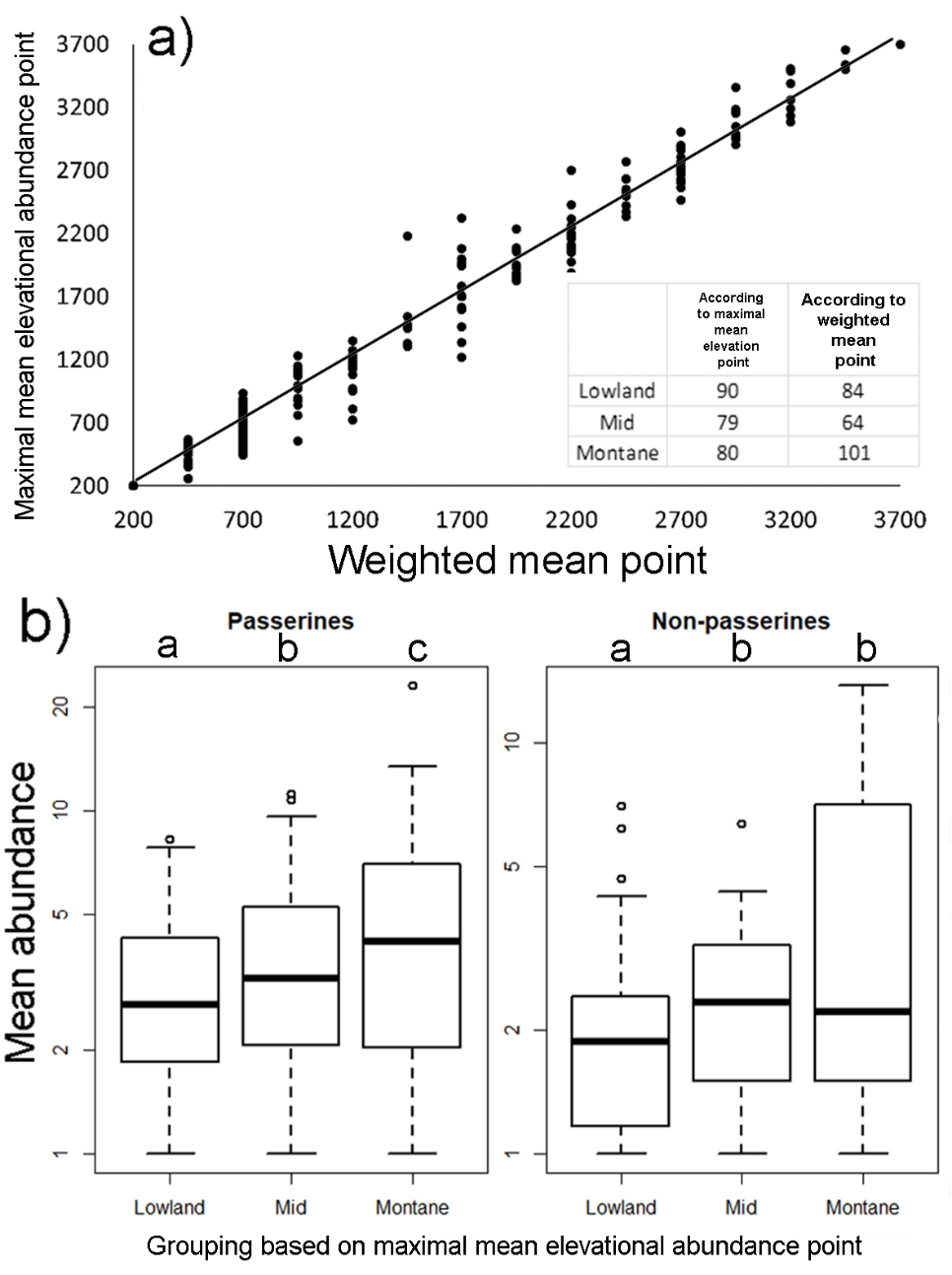
Figure S10. Correlation (Maximal mean elevational abundance point = 1.0198* Weighted mean point - 23.434, R² = 0.9745) between *weighted mean point* and *maximal mean elevational abundance* point (a) and number of species assigned to lowland, mid and montane group of species based on the weighted mean and maximal abundance points (insert in a). Passerine and non-passerine species are divided into three groups based on the position of their maximal mean elevational abundance point and their mean abundances and summarized across season (b). The pattern is also valid within season: Kruskal-Wallis - passerines in wet season (a) χ^2^ = 9.64, df = 2, N = 161, P < 0.008; in dry season (b) χ^2^ = 5.87, df = 2, N = 161, P < 0.05; non-passerines in wet season (c) χ^2^ = 4.75, df = 2, N = 88, P = 0.05; in dry season (d) χ^2^ = 6.04, df = 2, N = 88, P = 0.048. Lowland group = elevational maximal mean point up to 800m a.s.l., mid group = elevational maximal mean point between 801 and 1600m a.s.l., and montane group = elevational maximal mean point above 1600 m a.s.l.

Figure S11. Passerine (a) and non-passerine (b) birds divided into three groups based on the position of their *weighted mean point* of elevational distribution on Mt. Wilhelm, and the length of their elevational ranges. Kruskal-Wallis passerines (a): χ^2^ = 22.7, df = 2, N = 161, P < 0.001; non-passerines (b) χ^2^ = 10.8, df = 2, N = 88, P = 0.004. Significant differences between the groups of birds are denoted by different letters above the box-plots. Lowland group = elevational weighted mean point up to 800m a.s.l., mid group = elevational weighted mean point between 801 and 1600m a.s.l., and montane group = elevational weighted mean point above 1600 m a.s.l.

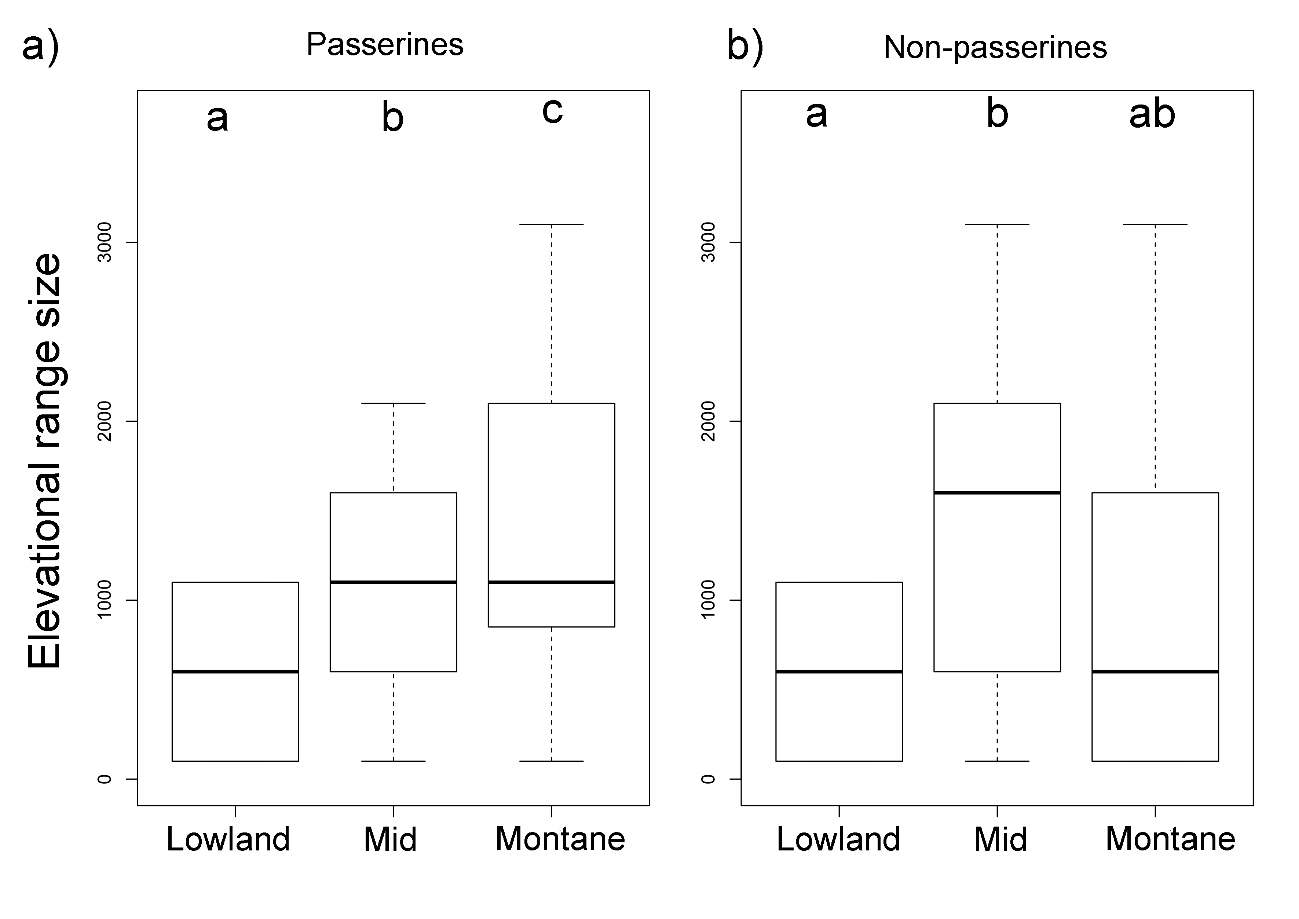

Figure S12. The body mass of passerine and non-passerine bird species and size of the geographical range they occupy. Passerines: F_1,159_ = 0.105, P=0.746; non-passerines: F_1,247_ = 1.24, P=0.268.

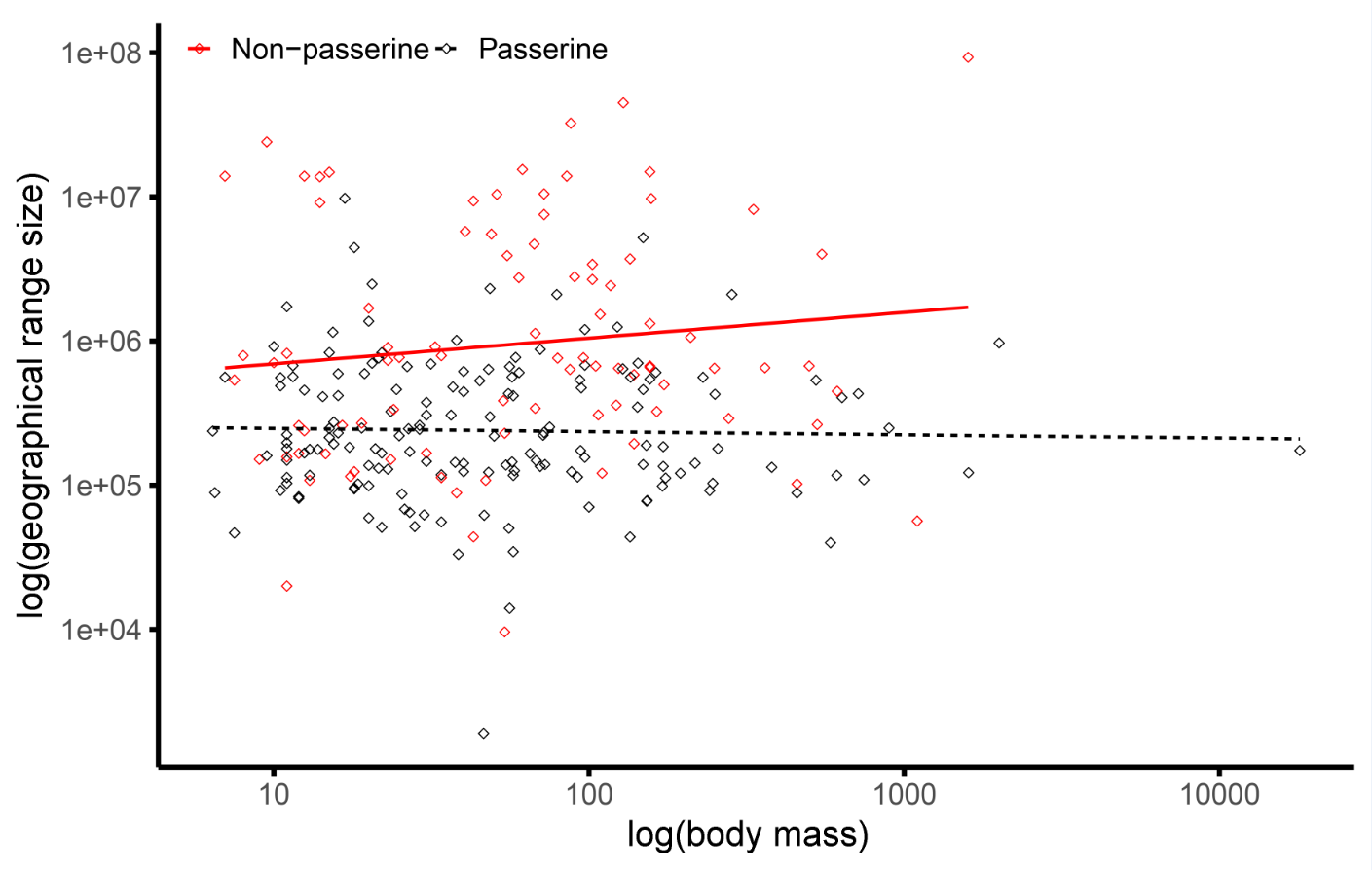

Table S1. List of bird species recorded during the point counts along the elevational gradient of Mt. Wilhelm in Papua New Guinea. Their *mean elevational abundance* (i.e. mean number of individuals recorded per 12.56 ha at each elevation where they were recorded) and *mean abundance* (i.e. across the range they occupied). Further, for each bird species the order is specified (PASS. for passerines and NON for non-passerines), the location its elevational mean-point and to which group of birds it was identified based on this weighted mean point (either lowland, mid-elevation or montane bird species). Finally, the last two columns show which feeding guild the species belong to and the size of their range (in km^2^). Feeding specialization was obtained from (Sam et al. 2019; Sam et al. 2017) and range are was obtained from Bird-Life International data zone.

| Scientific name | Mean elevational abundance at each elevation | | | | | | | | Mean abundance | Order | Weighted mean point | Group | Guild | Area |
| --- | --- | --- | --- | --- | --- | --- | --- | --- | --- | --- | --- | --- | --- | --- |
|  | 200 | 700 | 1200 | 1700 | 2200 | 2700 | 3200 | 3700 |  |  |  |  |  |  |
| *Acanthiza cinerea* |  |  |  | 1.83 | 1.56 | 5.77 | 1.67 |  | 2.706 | PASS. | 2450 | Mont. | In | 122000 |
| *Acanthiza murina* |  |  |  |  | 7 | 3.6 | 3.14 | 10.29 | 6.007 | PASS. | 2950 | Mont. | In | 83100 |
| *Accipiter fasciatus* |  |  |  |  | 2 |  |  |  | 2 | NON | 2200 | Mont. | Ca | 8000000 |
| *Accipiter meyerianus* |  |  |  |  | 1 |  |  |  | 1 | NON | 2200 | Mont. | Ca | 263000 |
| *Aegotheles albertisi* |  |  |  |  | 1 |  |  |  | 1 | NON | 2200 | Mont. | In | 88500 |
| *Aegotheles insignis* |  |  |  |  |  | 1.33 |  |  | 1.333 | NON | 2700 | Mont. | In | 166000 |
| *Aepypodius arfakianus* |  |  |  | 2.33 |  |  |  |  | 2.333 | NON | 1700 | Mid | Fr | 194000 |
| *Aerodramus hirundinaceus* |  |  |  | 2.5 |  |  |  |  | 2.5 | NON | 1700 | Mid | In | 584000 |
| *Ailuroedus buccoides* |  | 1.5 | 1 | 1 |  |  |  |  | 1.167 | PASS. | 1200 | Mid | Fr-In | 375000 |
| *Ailuroedus melanotis* |  |  |  |  | 1 |  |  |  | 1 | PASS. | 2200 | Mont. | Fr-In | 167000 |
| *Aleadryas rufinucha* |  |  |  | 1.4 | 4.4 | 2.33 | 2.73 | 1.8 | 2.532 | PASS. | 2700 | Mont. | In | 142000 |
| *Alisterus chloropterus* |  | 1 | 1 |  | 8.22 | 10 |  |  | 5.056 | NON | 1700 | Mid | Fr | 324000 |
| *Alopecoenas beccarii* |  |  | 1.33 | 1.67 |  |  |  |  | 1.5 | NON | 1450 | Mid | Fr | 167000 |
| *Alopecoenas jobiensis* |  |  | 1 |  | 4 |  |  |  | 2.5 | NON | 1700 | Mid | Fr | 647000 |
| *Amalocichla incerta* |  |  |  | 1 |  |  |  |  | 1 | PASS. | 1700 | Mont. | In | 144000 |
| *Amblyornis macgregoriae* |  |  |  | 1 |  | 1 | 2.63 |  | 1.542 | PASS. | 2450 | Mont. | Fr | 14000 |
| *Anthus gutturalis* |  |  |  |  |  |  | 10.8 | 16.07 | 13.436 | PASS. | 3450 | Mont. | In | 34600 |
| *Aplonis cantoroides* | 4.86 |  |  |  |  |  |  |  | 4.857 | PASS. | 200 | Low. | Fr-In | 831000 |
| *Aplonis metallica* | 10.71 |  |  |  |  |  |  |  | 10.714 | PASS. | 200 | Low. | Fr-In | 770000 |
| *Arses insularis* | 1.71 | 2.25 | 3.43 | 3 |  |  |  |  | 2.598 | PASS. | 950 | Mid | In | 249000 |
| *Artamus maximus* |  |  |  |  |  | 6.83 | 2 | 5 | 4.611 | PASS. | 3200 | Mont. | In | 249000 |
| *Astrapia stephaniae* |  |  |  |  | 1 | 2.14 | 10.17 | 2.6 | 3.977 | PASS. | 2950 | Mont. | Fr | 55600 |
| *Cacatua galerita* | 9 | 4.9 | 2.09 | 1 |  |  |  |  | 4.248 | NON | 950 | Mid | Fr | 4000000 |
| *Cacomantis castaneiventris* | 1 | 1 | 2 | 1.88 | 2.14 | 1 |  |  | 1.503 | NON | 1450 | Mid | In | 791000 |
| *Cacomantis flabelliformis* |  |  | 3 | 1.43 | 1.57 | 1.75 | 1.88 |  | 1.925 | NON | 2200 | Mont. | In | 2000000 |
| *Cacomantis leucolophus* | 1.4 | 1.25 | 3.22 |  |  |  |  |  | 1.957 | NON | 700 | Low. | In | 497000 |
| *Cacomantis variolosus* | 3 | 3.6 | 1.67 | 1.25 |  |  |  |  | 2.379 | NON | 950 | Mid | In | 4000000 |
| *Caligavis obscura* |  |  | 1 |  |  |  |  |  | 1 | PASS. | 1200 | Mid | Fr-In | 174000 |
| *Caligavis subfrenata* |  |  |  | 1.5 | 1 | 6.57 | 7.91 | 4.63 | 4.321 | PASS. | 2700 | Mont. | In-Ne | 133000 |
| *Campochaera sloetii* | 1.5 |  | 3.33 |  |  |  |  |  | 2.417 | PASS. | 700 | Low. | Fr-In | 230000 |
| *Caprimulgus macrurus* | 1 |  |  |  |  |  |  |  | 1 | NON | 200 | Low. | In | 6000000 |
| *Carterornis chrysomela* | 2.64 | 1.8 | 3.8 |  |  |  |  |  | 2.745 | PASS. | 700 | Low. | In | 641000 |
| *Casuarius bennetti* |  |  |  |  |  | 1 |  |  | 1 | NON | 2700 | Mont. | Fr | 359000 |
| *Centropus phasianinus* | 2.29 | 1 |  |  |  |  |  |  | 1.643 | NON | 450 | Low. | In | 3000000 |
| *Ceyx azureus* | 3 | 1.67 | 1.33 |  |  |  |  |  | 2 | NON | 700 | Low. | In | 3000000 |
| *Ceyx lepidus* | 5.11 | 6.43 | 7.58 |  |  |  |  |  | 6.374 | NON | 700 | Low. | In | 43800 |
| *Ceyx pusillus* | 1 |  |  |  |  |  |  |  | 1 | NON | 200 | Low. | In | 910000 |
| *Chaetorhynchus papuensis* | 1 | 1 | 3.22 | 2.17 |  |  |  |  | 1.847 | PASS. | 950 | Mid | In | 306000 |
| *Chalcophaps indica* | 1 | 1 |  |  |  |  |  |  | 1 | NON | 450 | Low. | Fr-In | 5000000 |
| *Chalcophaps stephani* |  | 1.67 | 1 |  |  |  |  |  | 1.333 | NON | 950 | Mid | Fr | 902000 |
| *Charmosyna josefinae* |  |  |  | 9.5 | 25 | 7 |  |  | 13.833 | NON | 2200 | Mont. | Ne | 151000 |
| *Charmosyna papou* |  |  |  | 4.4 | 8.86 | 10.57 | 12.67 | 3.8 | 8.059 | NON | 2700 | Mont. | Ne | 9600 |
| *Charmosyna placentis* | 2 | 2.5 |  |  |  |  |  |  | 2.25 | NON | 450 | Low. | Ne | 821000 |
| *Charmosyna rubronotata* | 2.33 |  |  |  |  |  |  |  | 2.333 | NON | 200 | Low. | Ne | 259000 |
| *Charmosyna wilhelminae* |  | 4 | 6.57 |  |  | 2.5 |  |  | 4.357 | NON | 1700 | Mid | Ne | 290000 |
| *Chlamydera lauterbachi* |  |  |  |  | 1 |  |  |  | 1 | PASS. | 2200 | Mont. | Fr-In | 124000 |
| *Chrysococcyx minutillus* | 1 |  |  |  |  |  |  |  | 1 | NON | 200 | Low. | In | 3000000 |
| *Chrysococcyx ruficollis* |  |  |  |  |  | 1.67 |  |  | 1.667 | NON | 2700 | Mont. | In | 151000 |
| *Cicinnurus regius* | 3.18 | 2.25 |  |  |  |  |  |  | 2.716 | PASS. | 450 | Low. | Fr-In | 480000 |
| *Cinnyris jugularis* | 3 | 4 | 5.29 | 7.38 |  |  |  |  | 4.915 | PASS. | 950 | Mid | In-Ne | 5000000 |
| *Clytoceyx rex* |  |  |  | 1.25 | 1 |  |  |  | 1.125 | NON | 1950 | Mont. | In | 341000 |
| *Clytomyias insignis* |  |  |  |  |  |  | 1 | 2 | 1.5 | PASS. | 3450 | Mont. | In | 139000 |
| *Cnemophilus loriae* |  |  |  | 1 | 1 | 1 | 1.5 |  | 1.125 | PASS. | 2450 | Mont. | Fr-In | 138000 |
| *Cnemophilus macgregorii* |  |  |  |  | 1.4 | 1.88 | 3 | 1.4 | 1.919 | PASS. | 2950 | Mont. | Fr | 43700 |
| *Collocalia esculenta* |  |  |  | 7 | 1.67 | 1 |  |  | 3.222 | NON | 2200 | Mont. | In | 3000000 |
| *Colluricincla megarhyncha* | 3.58 | 4.5 | 11.92 | 11.71 | 2.5 |  |  |  | 6.844 | PASS. | 1200 | Mid | In | 1000000 |
| *Columba vitiensis* |  |  |  |  | 1 | 2.33 |  |  | 1.667 | NON | 2450 | Mont. | Fr | 1000000 |
| *Coracina boyeri* | 1 | 10 | 1.67 |  |  |  |  |  | 4.222 | PASS. | 700 | Low. | Fr-In | 604000 |
| *Coracina caeruleogrisea* |  | 1 | 2.25 | 1.33 | 2 | 1.2 |  |  | 1.557 | PASS. | 1700 | Mont. | In | 405000 |
| *Coracina incerta* | 1 | 1.33 |  |  |  |  |  |  | 1.167 | PASS. | 450 | Low. | In | 348000 |
| *Coracina longicauda* |  |  |  | 1 |  | 2.67 |  |  | 1.833 | PASS. | 2200 | Mont. | In | 135000 |
| *Coracina melas* | 1.25 |  |  |  |  |  |  |  | 1.25 | PASS. | 200 | Low. | In | 593000 |
| *Coracina montana* |  | 1 | 4.33 | 5.73 | 1.5 | 1 |  |  | 2.712 | PASS. | 1700 | Mont. | Fr-In | 247000 |
| *Coracina papuensis* | 7.27 | 3.67 | 10 | 3.4 |  |  |  |  | 6.085 | PASS. | 950 | Mid | In | 4000000 |
| *Coracina schisticeps* |  |  |  |  | 1 |  |  |  | 1 | PASS. | 2200 | Mont. | Fr-In | 166000 |
| *Coracina tenuirostris* | 1 |  | 1.75 |  |  |  |  |  | 1.375 | PASS. | 700 | Low. | In | 2000000 |
| *Corvus tristis* | 4.88 | 3.67 | 3 | 3 |  |  |  |  | 3.635 | PASS. | 950 | Mid | Fr-In | 693000 |
| *Cracticus cassicus* | 7.83 |  |  |  |  |  |  |  | 7.833 | PASS. | 200 | Low. | Fr-In | 561000 |
| *Cracticus quoyi* | 1 |  |  |  |  |  |  |  | 1 | PASS. | 200 | Low. | In | 1000000 |
| *Crateroscelis murina* | 1 | 8.7 | 8.79 | 6.38 |  |  |  |  | 6.218 | PASS. | 950 | Mid | In | 237000 |
| *Crateroscelis nigrorufa* |  |  |  | 1.33 |  |  |  |  | 1.333 | PASS. | 1700 | Mont. | In | 114000 |
| *Crateroscelis robusta* |  | 4 |  | 3.44 | 5.33 | 6 | 9.46 | 9.36 | 6.266 | PASS. | 2200 | Mont. | In | 156000 |
| *Cyclopsitta diophthalma* | 1.5 | 4 | 8.54 | 2.8 |  |  |  |  | 4.21 | NON | 950 | Mid | Fr | 448000 |
| *Cyclopsitta gulielmitertii* | 2.25 |  | 1.5 |  |  |  |  |  | 1.875 | NON | 700 | Low. | Fr | 102000 |
| *Dacelo gaudichaud* | 10.91 | 1.5 |  |  |  |  |  |  | 6.205 | NON | 450 | Low. | In | 671000 |
| *Daphoenositta miranda* |  |  |  |  |  | 2.5 | 1.71 | 1.25 | 1.821 | PASS. | 3200 | Mont. | In | 39900 |
| *Dicaeum geelvinkianum* | 3.2 | 7.1 | 6.93 | 12.31 | 5.85 |  |  |  | 7.076 | PASS. | 1200 | Mid | Fr | 535000 |
| *Dicrurus bracteatus* | 6.17 | 3.33 |  |  |  |  |  |  | 4.75 | PASS. | 450 | Low. | In | 2000000 |
| *Diphyllodes magnificus* |  | 3.8 | 4.43 | 2.33 |  |  |  |  | 3.521 | PASS. | 1200 | Mid | Fr-In | 112000 |
| *Ducula chalconota* |  |  |  | 1.33 | 3.43 | 1 |  |  | 1.921 | NON | 2200 | Mont. | Fr | 165000 |
| *Ducula pinon* | 1.43 | 1.5 |  |  |  |  |  |  | 1.464 | NON | 450 | Low. | Fr | 635000 |
| *Ducula rufigaster* | 1 |  |  |  |  |  |  |  | 1 | NON | 200 | Low. | Fr | 671000 |
| *Ducula zoeae* | 7 | 2.43 | 4.62 |  |  |  |  |  | 4.681 | NON | 700 | Low. | Fr | 707000 |
| *Eclectus roratus* | 7.08 | 3.78 | 1 |  |  |  |  |  | 3.954 | NON | 700 | Low. | Fr | 2000000 |
| *Epimachus fastosus* |  |  | 1 | 1.5 | 2.67 | 4.09 |  |  | 2.314 | PASS. | 1950 | Mont. | Fr-In | 78200 |
| *Epimachus meyeri* |  |  |  | 2 | 3.5 | 8.77 | 4.8 |  | 4.767 | PASS. | 2450 | Mont. | Fr-In | 135000 |
| *Erythropitta erythrogaster* | 1.6 | 3.2 |  |  |  |  |  |  | 2.4 | PASS. | 450 | Low. | In | 1000000 |
| *Erythrura trichroa* |  |  |  | 4 | 2.33 | 1.6 | 2 | 7 | 3.387 | PASS. | 2700 | Mont. |  | 875000 |
| *Eudynamys scolopaceus* | 2.4 | 2.5 |  |  |  |  |  |  | 2.45 | NON | 450 | Low. | Fr-In | 10000000 |
| *Eugerygone rubra* |  |  |  | 1 | 2.38 | 3.75 | 1.67 | 2.11 | 2.181 | PASS. | 2700 | Mont. | In | 121000 |
| *Eulacestoma nigropectus* |  |  |  |  |  | 2.75 |  |  | 2.75 | PASS. | 2700 | Mont. | In | 88700 |
| *Eurystomus orientalis* | 1.17 | 3 |  |  |  |  |  |  | 2.083 | NON | 450 | Low. | In | 10000000 |
| *Garritornis isidorei* | 2.5 |  |  |  |  |  |  |  | 2.5 | PASS. | 200 | Low. | In | 561000 |
| *Geoffroyus geoffroyi* | 2.8 |  |  |  |  |  |  |  | 2.8 | NON | 200 | Low. | Fr | 793000 |
| *Geoffroyus simplex* | 1 |  |  |  |  |  |  |  | 1 | NON | 200 | Low. | Fr | 238000 |
| *Gerygone chloronota* | 1.5 | 2.67 | 2.38 |  |  |  |  |  | 2.181 | PASS. | 700 | Low. | In | 1000000 |
| *Gerygone chrysogaster* | 2.5 | 3.78 |  |  |  |  |  |  | 3.139 | PASS. | 450 | Low. | In | 544000 |
| *Gerygone palpebrosa* | 1.67 |  | 1.8 |  |  |  |  |  | 1.733 | PASS. | 700 | Low. | In | 969000 |
| *Gerygone ruficollis* |  |  |  | 6.18 | 4.38 | 7.75 | 3.17 | 1.33 | 4.561 | PASS. | 2700 | Mont. | In | 103000 |
| *Grallina bruijnii* |  |  | 3 | 1 |  |  |  |  | 2 | PASS. | 1450 | Mid | In | 260000 |
| *Gymnophaps albertisii* |  | 4 |  | 3.78 | 14.67 | 11.15 | 1 | 2 | 6.1 | NON | 2200 | Mont. | Fr | 536000 |
| *Harpyopsis novaeguineae* |  |  | 1 | 2 |  | 1 |  |  | 1.333 | NON | 1950 | Mont. | Ca | 734000 |
| *Henicophaps albifrons* | 1 | 1 | 1 |  |  |  |  |  | 1 | NON | 700 | Low. | Fr | 769000 |
| *Heteromyias albispecularis* |  |  |  | 3.43 | 1 | 1 |  |  | 1.81 | PASS. | 2200 | Mont. | In | 123000 |
| *Ifrita kowaldi* |  |  |  | 2 | 2 | 9.6 | 7.08 | 3.29 | 4.794 | PASS. | 2700 | Mont. | In | 91900 |
| *Lalage atrovirens* | 1 |  |  |  |  |  |  |  | 1 | PASS. | 200 | Low. | Fr-In | 306000 |
| *Leptocoma sericea* | 7.83 | 1.5 | 3.13 |  |  |  |  |  | 4.153 | PASS. | 700 | Low. | In-Ne | 915000 |
| *Loboparadisea sericea* |  |  |  | 1.5 |  | 1 |  |  | 1.25 | PASS. | 2200 | Mont. | Fr | 174000 |
| *Lonchura spectabilis* |  |  |  | 1 | 3.33 |  |  |  | 2.167 | PASS. | 1950 | Mont. | Gr | 214000 |
| *Lonchura tristissima* | 4 |  |  |  |  |  |  |  | 4 | PASS. | 200 | Low. | Gr | 560000 |
| *Lophorina superba* |  |  |  | 3.57 |  |  |  |  | 3.571 | PASS. | 1700 | Mont. | Fr-In | 160000 |
| *Loriculus aurantiifrons* | 2.22 |  |  |  |  |  |  |  | 2.222 | NON | 200 | Low. | Ne | 20000 |
| *Lorius lory* | 5.43 | 12.11 | 3.55 |  |  |  |  |  | 7.028 | NON | 700 | Low. | Ne | 10000000 |
| *Machaerirhynchus flaviventer* | 1.13 | 2 | 3.85 | 1 |  |  |  |  | 1.993 | PASS. | 950 | Mid | In | 702000 |
| *Machaerirhynchus nigripectus* |  |  | 6 | 4.5 | 2 | 3 | 1.33 |  | 3.367 | PASS. | 2200 | Mont. | In | 219000 |
| *Macropygia amboinensis* | 3.9 | 2 | 4 | 4.27 | 3.22 |  |  |  | 3.479 | NON | 1200 | Mid | Fr | 1000000 |
| *Macropygia nigrirostris* |  | 1 |  | 5 | 12 | 2.86 |  |  | 5.214 | NON | 1700 | Mid | Fr | 647000 |
| *Malurus alboscapulatus* |  |  |  | 2.5 | 6 |  |  |  | 4.25 | PASS. | 1950 | Mont. | In | 431000 |
| *Manucodia chalybatus* |  |  | 1.4 |  |  |  |  |  | 1.4 | PASS. | 1200 | Mid | Fr | 81000 |
| *Megalurus macrurus* |  |  |  | 2 |  |  |  |  | 2 | PASS. | 1700 | Mont. | In | 2000000 |
| *Megapodius decollatus* | 1 | 1.67 |  |  |  |  |  |  | 1.333 | NON | 450 | Low. | Fr-In | 10000000 |
| *Melampitta lugubris* |  |  |  |  |  | 3 | 2.14 | 4 | 3.048 | PASS. | 3200 | Mont. | In | 59300 |
| *Melanocharis longicauda* |  |  |  | 1 |  | 1 |  |  | 1 | PASS. | 2200 | Mont. | Fr-In | 94300 |
| *Melanocharis nigra* | 5.5 | 12.33 | 6.36 | 6 | 1 |  |  |  | 6.238 | PASS. | 1200 | Mid | Fr-In | 461000 |
| *Melanocharis striativentris* |  |  |  | 2.78 |  | 1.5 |  |  | 2.139 | PASS. | 2200 | Mont. | Fr | 86800 |
| *Melanocharis versteri* |  |  |  | 5 | 7.92 | 5.69 | 5.54 | 4 | 5.631 | PASS. | 2700 | Mont. | Fr-In | 145000 |
| *Melanorectes nigrescens* |  |  |  | 2.57 | 2.8 | 2 |  |  | 2.457 | PASS. | 2200 | Mont. | In | 126000 |
| *Melidectes belfordi* |  |  |  | 10 | 22.43 | 30.57 | 39.08 | 13.91 | 23.197 | PASS. | 2700 | Mont. | In-Ne | 124000 |
| *Melidectes fuscus* |  |  |  |  | 3.86 | 1.71 | 6.08 | 18.85 | 7.624 | PASS. | 2950 | Mont. | In-Ne | 70500 |
| *Melidectes princeps* |  |  |  |  |  |  | 1 | 9.64 | 5.318 | PASS. | 3450 | Mont. | In-Ne | 1900 |
| *Melidectes rufocrissalis* |  |  |  | 9.44 | 1 | 1.5 |  |  | 3.981 | PASS. | 2200 | Mont. | Fr-In | 64700 |
| *Melidectes torquatus* |  |  | 2.5 | 4.73 | 1 |  |  |  | 2.742 | PASS. | 1700 | Mont. | Fr-In | 95800 |
| *Melidora macrorrhina* | 1 | 2 |  |  |  |  |  |  | 1.5 | NON | 450 | Low. | In | 108000 |
| *Melilestes megarhynchus* | 3 | 4.1 | 2.83 | 2.13 | 1 |  |  |  | 2.612 | PASS. | 1200 | Mid | In-Ne | 562000 |
| *Meliphaga analoga* | 18.58 | 8.4 | 9.27 | 4.13 | 1 |  |  |  | 8.276 | PASS. | 1200 | Mid | In-Ne | 636000 |
| *Meliphaga aruensis* | 1.83 | 1.5 | 3.25 |  |  |  |  |  | 2.194 | PASS. | 700 | Low. | Fr-In | 664000 |
| *Meliphaga montana* |  |  | 3.88 |  |  |  |  |  | 3.875 | PASS. | 1200 | Mid | Fr-In | 118000 |
| *Meliphaga orientalis* |  |  |  | 8.5 | 1.25 | 1.25 |  |  | 3.667 | PASS. | 2200 | Mont. | In-Ne | 193000 |
| *Melipotes fumigatus* |  |  | 3.5 | 4.17 | 3.5 | 5.5 | 8.33 | 4.89 | 4.981 | PASS. | 2450 | Mont. | Fr-In | 149000 |
| *Merops ornatus* | 2 |  |  |  |  |  |  |  | 2 | NON | 200 | Low. | In | 13760000 |
| *Microdynamis parva* | 2 |  |  |  |  |  |  |  | 2 | NON | 200 | Low. | Fr | 9360000 |
| *Microeca flavovirescens* | 2.63 | 4.57 | 4.22 |  |  |  |  |  | 3.806 | PASS. | 700 | Low. | In | 675000 |
| *Microeca griseoceps* |  |  | 1 |  |  |  |  |  | 1 | PASS. | 1200 | Mid | In | 189000 |
| *Microeca papuana* |  |  |  | 2.23 | 6.7 | 5.54 |  |  | 4.823 | PASS. | 2200 | Mont. | In | 142000 |
| *Micropsitta bruijnii* |  |  | 3 |  |  |  |  |  | 3 | NON | 1200 | Mid | In-Ne | 269000 |
| *Micropsitta pusio* | 6.57 | 6.29 | 5 |  |  |  |  |  | 5.952 | NON | 700 | Low. | In-Ne | 9120000 |
| *Mino anais* | 1 | 1 |  |  |  |  |  |  | 1 | PASS. | 450 | Low. | Fr | 411000 |
| *Mino dumontii* | 4.43 | 2.38 |  |  |  |  |  |  | 3.402 | PASS. | 450 | Low. | Fr-In | 701000 |
| *Monachella muelleriana* | 1.67 |  |  |  |  |  |  |  | 1.667 | PASS. | 200 | Low. | In | 418000 |
| *Monarcha frater* |  |  | 2.67 |  |  |  |  |  | 2.667 | PASS. | 1200 | Mid | In | 179000 |
| *Monarcha rubiensis* | 1.33 |  |  |  |  |  |  |  | 1.333 | PASS. | 200 | Low. | In | 244000 |
| *Myiagra alecto* | 2.56 | 2 | 1 |  |  |  |  |  | 1.852 | PASS. | 700 | Low. | In | 1000000 |
| *Myzomela rosenbergii* |  |  | 1.5 | 11 | 28.14 | 4.64 | 5.62 | 4.3 | 9.199 | PASS. | 2450 | Mont. | In-Ne | 177000 |
| *Neopsittacus musschenbroekii* |  |  | 6.5 | 5.63 | 2.33 | 2.67 | 1.5 |  | 3.725 | NON | 2200 | Mont. | Ne | 229000 |
| *Neopsittacus pullicauda* |  |  |  | 6.13 | 5.2 | 10.56 | 11.18 | 12 | 9.012 | NON | 2700 | Mont. | Ne | 113000 |
| *Oedistoma iliolophus* |  | 6.67 | 9.23 | 2.44 |  |  |  |  | 6.114 | PASS. | 1200 | Mid | In | 557000 |
| *Oreocharis arfaki* |  |  |  | 2.91 | 3.25 | 5 | 2 | 2.5 | 3.132 | PASS. | 2700 | Mont. | Fr | 50200 |
| *Oreopsittacus arfaki* |  |  |  |  | 3.43 | 11.43 | 20.25 | 16.22 | 12.832 | NON | 2950 | Mont. | Ne | 108000 |
| *Oreostruthus fuliginosus* |  |  |  |  |  |  |  | 5.8 | 5.8 | PASS. | 3700 | Mont. | Fr-In | 51000 |
| *Oriolus szalayi* | 5.14 |  |  |  |  |  |  |  | 5.143 | PASS. | 200 | Low. | Fr-In | 680000 |
| *Ornorectes cristatus* |  |  | 2.5 |  |  |  |  |  | 2.5 | PASS. | 1200 | Mid | In | 88200 |
| *Otidiphaps nobilis* |  |  | 1 |  |  |  |  |  | 1 | NON | 1200 | Mid | Fr | 260000 |
| *Pachycare flavogriseum* |  |  | 1.33 | 1.33 |  |  |  |  | 1.333 | PASS. | 1450 | Mid | In | 171000 |
| *Pachycephala hyperythra* | 3 | 1.17 | 9.73 | 5.29 |  |  |  |  | 4.795 | PASS. | 950 | Mid | In | 99100 |
| *Pachycephala modesta* |  |  |  |  |  | 2.25 | 3 |  | 2.625 | PASS. | 2950 | Mont. | In | 68100 |
| *Pachycephala monacha* |  | 1 |  |  |  |  |  |  | 1 | PASS. | 700 | Low. | In | 33200 |
| *Pachycephala schlegelii* |  |  |  | 6.9 | 9.29 | 15.64 | 6.17 | 4.3 | 8.459 | PASS. | 2700 | Mont. | In | 129000 |
| *Pachycephala simplex* |  | 3 | 3.5 |  |  |  |  |  | 3.25 | PASS. | 950 | Mid | In | 829000 |
| *Pachycephala soror* |  | 3.5 | 7.2 | 4.27 | 2.22 | 1.5 |  |  | 3.739 | PASS. | 1700 | Mont. | In | 220000 |
| *Pachycephalopsis poliosoma* |  |  | 7.83 | 2.83 |  |  |  |  | 5.333 | PASS. | 1450 | Mid | In | 185000 |
| *Paradigalla brevicauda* |  |  |  |  | 1 |  |  |  | 1 | PASS. | 2200 | Mont. | Fr-In | 91700 |
| *Paradisaea minor* | 8.5 | 9.6 | 15.39 |  |  |  |  |  | 11.162 | PASS. | 700 | Low. | Fr-In | 298000 |
| *Paramythia montium* |  |  |  |  |  | 3.58 | 8.21 | 27.23 | 13.009 | PASS. | 3200 | Mont. | Fr | 62200 |
| *Peltops blainvillii* | 2.44 | 1.29 |  |  |  |  |  |  | 1.865 | PASS. | 450 | Low. | In | 530000 |
| *Peltops montanus* |  | 1.5 |  | 4 | 1 | 3.67 |  |  | 2.542 | PASS. | 1700 | Mont. | In | 324000 |
| *Peneothello bimaculata* |  | 6.83 | 6.86 | 8.56 |  |  |  |  | 7.415 | PASS. | 1200 | Mid | In | 51600 |
| *Peneothello cyanus* |  |  |  | 14.39 | 17.5 | 5 |  |  | 12.295 | PASS. | 2200 | Mont. | In | 167000 |
| *Peneothello sigillata* |  |  |  |  |  | 11.25 | 9.92 | 10.42 | 10.53 | PASS. | 3200 | Mont. | In | 77400 |
| *Philemon buceroides* | 9.82 | 1.33 |  |  |  |  |  |  | 5.576 | PASS. | 450 | Low. | In-Ne | 432000 |
| *Philemon meyeri* | 7.08 | 3.5 | 2.17 |  |  |  |  |  | 4.25 | PASS. | 700 | Low. | In-Ne | 46600 |
| *Phylloscopus maforensis* |  |  | 2.33 | 4.27 | 1 |  |  |  | 2.535 | PASS. | 1700 | Mont. | In | 473000 |
| *Pitohui dichrous* |  | 6.88 | 15.07 | 5.64 |  |  |  |  | 9.196 | PASS. | 1200 | Mid | Fr-In | 222000 |
| *Pitohui kirhocephalus* | 3.4 | 8.6 | 8.2 |  |  |  |  |  | 6.733 | PASS. | 700 | Low. | In | 538000 |
| *Pitta sordida* | 2 | 2 |  |  |  |  |  |  | 2 | PASS. | 450 | Low. | In | 2000000 |
| *Podargus ocellatus* |  |  |  | 1 |  |  |  |  | 1 | NON | 1700 | Mid | In | 761000 |
| *Poecilodryas albonotata* |  |  |  |  | 1.25 | 1 | 1.2 |  | 1.15 | PASS. | 2700 | Mont. | In | 117000 |
| *Poecilodryas hypoleuca* | 3.75 | 6 | 3.17 |  |  |  |  |  | 4.306 | PASS. | 700 | Low. | In | 417000 |
| *Probosciger aterrimus* | 3.36 | 2.38 | 1.6 |  |  |  |  |  | 2.446 | NON | 700 | Low. | Fr | 14880000 |
| *Pseudeos fuscata* | 3.11 |  |  | 5.75 | 20.27 | 16.43 |  |  | 11.391 | NON | 1450 | Mid | Fr-In | 766000 |
| *Pseudorectes ferrugineus* | 7.83 |  | 4 |  |  |  |  |  | 5.917 | PASS. | 700 | Low. | Fr-In | 615000 |
| *Psittacella brehmii* |  |  |  |  | 1 | 2 |  |  | 1.5 | NON | 2450 | Mont. | Fr | 124000 |
| *Psittacella picta* |  |  |  |  |  | 1.25 | 2 | 4 | 2.417 | NON | 3200 | Mont. | Fr | 56400 |
| *Psittaculirostris edwardsii* | 3 | 3 | 2.8 |  |  |  |  |  | 2.933 | NON | 700 | Low. | Fr | 1320000 |
| *Psitteuteles goldiei* |  |  |  |  |  | 13 | 13 |  | 13 | NON | 2950 | Mont. | Ne | 307000 |
| *Psittrichas fulgidus* | 2 |  |  |  |  |  |  |  | 2 | NON | 200 | Low. | Fr | 5512000 |
| *Pteridophora alberti* |  |  |  |  |  | 1 |  |  | 1 | PASS. | 2700 | Mont. | Fr-In | 109000 |
| *Ptilinopus coronulatus* | 1.5 | 1 | 2.25 | 4.6 |  |  |  |  | 2.338 | NON | 950 | Mid | Fr | 670000 |
| *Ptilinopus iozonus* | 5.33 |  |  |  |  |  |  |  | 5.333 | NON | 200 | Low. | Fr | 10400000 |
| *Ptilinopus magnificus* | 2.7 |  | 2.2 |  |  |  |  |  | 2.45 | NON | 700 | Low. | Fr | 32400000 |
| *Ptilinopus ornatus* |  | 2.5 |  | 1 | 1.25 |  |  |  | 1.583 | NON | 1450 | Mid | Fr | 385000 |
| *Ptilinopus perlatus* | 1 | 1.5 |  |  |  |  |  |  | 1.25 | NON | 450 | Low. | Fr | 10480000 |
| *Ptilinopus pulchellus* | 1.88 | 1.33 | 1.5 |  |  |  |  |  | 1.569 | NON | 700 | Low. | Fr | 7536000 |
| *Ptilinopus rivoli* |  |  |  | 4 | 4.75 | 3.75 | 3.75 |  | 4.063 | NON | 2450 | Mont. | Fr | 335000 |
| *Ptilinopus superbus* | 1.67 | 1.33 | 4.14 |  | 2 |  |  |  | 2.286 | NON | 1200 | Mid | Fr | 2000000 |
| *Ptiloprora guisei* |  |  |  | 3 | 6.5 | 5 | 1.8 |  | 4.075 | PASS. | 2450 | Mont. | Fr-In | 61900 |
| *Ptiloprora meekiana* |  |  |  | 2 |  |  |  |  | 2 | PASS. | 1700 | Mont. | In | 139000 |
| *Ptiloprora perstriata* |  |  |  |  | 3.5 | 18.86 | 14.79 | 6.2 | 10.836 | PASS. | 2950 | Mont. | In | 102000 |
| *Ptiloris magnificus* |  | 2 | 8.31 |  |  |  |  |  | 5.154 | PASS. | 950 | Mid | Fr-In | 605000 |
| *Ptilorrhoa caerulescens* | 2 | 1.83 | 2 |  |  |  |  |  | 1.944 | PASS. | 700 | Low. | In | 427000 |
| *Ptilorrhoa castanonota* |  |  | 2.33 |  |  |  |  |  | 2.333 | PASS. | 1200 | Mid | In | 246000 |
| *Ptilorrhoa leucosticta* |  |  |  | 1.4 | 1.5 | 2 |  |  | 1.633 | PASS. | 2200 | Mont. | In | 232000 |
| *Pycnopygius ixoides* | 1.33 | 1 | 4.5 |  |  |  |  |  | 2.278 | PASS. | 700 | Low. | Fr | 460000 |
| *Rallicula forbesi* |  |  |  |  |  | 1.5 |  |  | 1.5 | NON | 2700 | Mont. | In | 121000 |
| *Reinwardtoena reinwardti* | 1 | 1.5 | 2 | 1.33 | 1.63 | 1.8 | 1.5 |  | 1.537 | NON | 1700 | Mid | Fr | 656000 |
| *Rhagologus leucostigma* |  |  |  | 3.8 | 2.7 | 2.25 |  |  | 2.917 | PASS. | 2200 | Mont. | Fr-In | 146000 |
| *Rhipidura albolimbata* |  |  |  | 11.07 | 12.33 | 12.29 | 8.46 | 6 | 10.03 | PASS. | 2700 | Mont. | In | 148000 |
| *Rhipidura atra* |  | 1.5 | 3.43 | 10.79 | 7.29 | 6.9 |  |  | 5.98 | PASS. | 1700 | Mont. | In | 179000 |
| *Rhipidura brachyrhyncha* |  |  |  |  | 4.13 | 10.67 | 6.62 | 3.88 | 6.321 | PASS. | 2950 | Mont. | In | 131000 |
| *Rhipidura hyperythra* |  | 4 |  |  |  |  |  |  | 4 | PASS. | 700 | Low. | In | 456000 |
| *Rhipidura leucothorax* | 3.83 | 1.5 | 1 |  |  |  |  |  | 2.111 | PASS. | 700 | Low. | In | 565000 |
| *Rhipidura rufidorsa* |  | 3 |  |  |  |  |  |  | 3 | PASS. | 700 | Low. | In | 488000 |
| *Rhipidura rufiventris* | 3.67 | 3.88 | 9.08 |  |  |  |  |  | 5.54 | PASS. | 700 | Low. | In | 2000000 |
| *Rhipidura threnothorax* | 6.75 | 3.75 | 6.13 |  | 1.5 |  |  |  | 4.531 | PASS. | 1200 | Mid | In | 594000 |
| *Rhyticeros plicatus* | 7.58 | 3.67 | 4.45 |  |  |  |  |  | 5.235 | NON | 700 | Low. | Fr | 24000000 |
| *Saxicola caprata* |  |  |  | 2 | 1 |  |  |  | 1.5 | PASS. | 1950 | Mont. | In | 10000000 |
| *Scolopax rosenbergii* |  |  |  |  |  | 1 |  |  | 1 | NON | 2700 | Mont. | In | 115000 |
| *Scythrops novaehollandiae* | 2 |  |  |  |  |  |  |  | 2 | NON | 200 | Low. | Fr-In | 92800000 |
| *Sericornis arfakianus* |  |  | 3 |  |  |  |  |  | 3 | PASS. | 1200 | Mid | In | 177000 |
| *Sericornis nouhuysi* |  |  |  | 5.17 | 12.39 | 17.79 | 12 | 6.33 | 10.734 | PASS. | 2700 | Mont. | In | 98600 |
| *Sericornis papuensis* |  |  |  | 7.67 | 7 | 18.25 | 6.36 |  | 9.82 | PASS. | 2450 | Mont. | In | 117000 |
| *Sericornis perspicillatus* |  |  |  | 15.14 | 18.83 | 4.86 |  |  | 12.944 | PASS. | 2200 | Mont. | In | 117000 |
| *Sericornis spilodera* |  | 4.5 | 5.33 | 2 |  | 1 |  |  | 3.208 | PASS. | 1700 | Mont. | In | 274000 |
| *Syma megarhyncha* |  |  | 2.71 | 2.67 | 2.33 | 2 |  |  | 2.429 | NON | 1950 | Mont. | In | 157000 |
| *Syma torotoro* | 1 | 2.75 |  |  |  |  |  |  | 1.875 | NON | 450 | Low. | In | 14800000 |
| *Symposiachrus axillaris* |  |  | 3.8 | 5 | 2.08 | 3 |  |  | 3.471 | PASS. | 1950 | Mont. | In | 113000 |
| *Symposiachrus guttula* | 2.6 | 3.75 | 1 |  |  |  |  |  | 2.45 | PASS. | 700 | Low. | In | 664000 |
| *Symposiachrus manadensis* | 4.71 |  |  |  |  |  |  |  | 4.714 | PASS. | 200 | Low. | In | 445000 |
| *Talegalla jobiensis* | 2.78 | 1.6 | 1.17 |  |  |  |  |  | 1.848 | NON | 700 | Low. | Fr-In | 4000000 |
| *Tanysiptera galatea* | 2.09 | 1.4 |  |  |  |  |  |  | 1.745 | NON | 450 | Low. | In | 15440000 |
| *Timeliopsis fulvigula* |  |  |  | 3.6 |  |  |  |  | 3.6 | PASS. | 1700 | Mont. | In | 137000 |
| *Toxorhamphus novaeguineae* | 7.92 | 8 | 9.29 |  |  |  |  |  | 8.401 | PASS. | 700 | Low. | In-Ne | 197000 |
| *Toxorhamphus poliopterus* |  |  | 6 | 12.5 | 10.29 |  |  |  | 9.595 | PASS. | 1700 | Mont. | In-Ne | 179000 |
| *Tregellasia leucops* |  | 1.5 | 5.2 |  |  |  |  |  | 3.35 | PASS. | 950 | Mid | In | 183000 |
| *Trichoglossus haematodus* | 13.42 | 7.4 | 4.9 |  |  |  |  |  | 8.572 | NON | 700 | Low. | Ne | 44880000 |
| *Trugon terrestris* |  |  |  | 1 | 1 |  |  |  | 1 | NON | 1950 | Mont. | Fr | 652000 |
| *Turdus poliocephalus* |  |  |  |  |  | 1.5 | 7.67 | 15.86 | 8.341 | PASS. | 3200 | Mont. | In | 253000 |
| *Xanthotis flaviventer* |  | 5.86 | 3.2 |  |  |  |  |  | 4.529 | PASS. | 950 | Mid | In | 762000 |
| *Zosterops minor* | 2 | 7.2 | 4.33 |  |  |  |  |  | 4.511 | PASS. | 700 | Low. | In | 224000 |
| *Zosterops novaeguineae* |  | 2 |  | 3.92 | 5.64 | 3 |  |  | 3.64 | PASS. | 1700 | Mont. | In | 103000 |

Sam, K., B. Koane, D. C. Bardos, S. Jeppy, and V. Novotny. 2019. Species richness of birds along a complete rain forest elevational gradient in the tropics: Habitat complexity and food resources matter. Journal of Biogeography **46**:279-290.
